# A new analysis of resting state connectivity and graph theory reveals distinctive short-term modulations due to whisker stimulation in rats

**DOI:** 10.1101/223057

**Authors:** Silke Kreitz, Benito de Celis Alonso, Michael Uder, Andreas Hess

## Abstract

Resting state (RS) connectivity has been increasingly studied in healthy and diseased brains in humans and animals. This paper presents a new method to analyze RS data from fMRI that combines multiple seed correlation analysis with graph-theory (MSRA). We characterize and evaluate this new method in relation to two other graph-theoretical methods and ICA. The graph-theoretical methods calculate cross-correlations of regional average time-courses, one using seed regions of the same size (SRCC) and the other using whole brain structure regions (RCCA). We evaluated the reproducibility, power, and capacity of these methods to characterize short term RS modulation to unilateral physiological whisker stimulation in rats. Graph-theoretical networks found with the MSRA approach were highly reproducible, and their communities showed large overlaps with ICA components. Additionally, MSRA was the only one of all tested methods that had the power to detect significant RS modulations induced by whisker stimulation that are controlled by family-wise error rate (FWE). Compared to the reduced resting state network connectivity during task performance, these modulations implied decreased connectivity strength in the bilateral sensorimotor and entorhinal cortex. Additionally, the contralateral ventromedial thalamus (part of the barrel field related lemniscal pathway) and the hypothalamus showed reduced connectivity. Enhanced connectivity was observed in the amygdala, especially the contralateral basolateral amygdala (involved in emotional learning processes). In conclusion, MSRA is a powerful analytical approach that can reliably detect tiny modulations of RS connectivity. It shows a great promise as a method for studying RS dynamics in healthy and pathological conditions.

## 1. Introduction

Since Biswal et al. (1995) first described intrinsic functional connectivity in the human brain during rest by functional MRI (fMRI), the so called resting state (RS) connectivity has been studied intensively and several large scale neural networks have been detected [see Raichle (2011) for review]. Despite the influence of individual and instantaneous factors such as mood, physiological and cognitive states, RS networks are remarkably robust and stable over time (Braun et al., 2012; Du et al., 2015; Zuo and Xing, 2014) and species (Gozzi and Schwarz, 2016; Lu et al., 2012; Sierakowiak et al., 2015). Thus, they seem to exhibit an evolutionary conserved and fundamental phenomenon of mammalian brain function. However, its biological relevance is still not fully understood. One hypothesis interprets human RS connectivity as a correlate of introspective mental processes (including such processes as mind wandering) that influences behavioral responses to future events (Rosazza and Minati, 2011). Other researchers emphasize the relation of RS topography and strength to the history of network activation and thus support the Hebbian-like Fire-Wire-hypothesis: regions that are co-activated during task performance develop stronger coherence at rest (Corbetta, 2010; Harmelech et al., 2013). Regardless of other interpretations, it has been widely accepted that resting state networks are dynamic in nature (Deco and Corbetta, 2011). They are modulated by prior task activation, which supports the hypothesis that RS functional connectivity plays a role in learning processes and memory consolidation (Albert et al., 2009; Tambini et al., 2010).

Short-term modulations of RS connectivity have also been detected in several human studies after performance of a motor task (Mary et al., 2016; Muraskin et al., 2016; Tung et al., 2013), but to the best of our knowledge, studies using sensory stimuli as RS modulators or animal models have not been published. One reason might be that we lack adequate analytical tools to detect these tiny modulations.

Therefore, our goal is to develop a method that is capable of examining RS connectivity during sensorimotor stimulation of the barrel field in the rat. For that purpose we evaluated three graph-theoretical resting state analysis methods in comparison to ICA to determine which analytical methods can best detect significant and reliable short term resting state modulations. The barrel field model and its underlying functional and anatomical pathways have been intensively investigated with various neurobiological techniques including fMRI (Diamond et al., 2008; Grinvald et al., 1986; Hess et al., 2000; Sachdev et al., 2003; Yang et al., 1997). We used unilateral stimulation of a small set of whiskers after trimming the remaining ones. Unilateral whisker activation usually indicates food or foe; thus the animal reacts with an immediate behavioral and emotional response (Marshall et al., 1971; Prescott, 2011). When compared to the more natural stimulation of all whiskers, trimming and stimulation of the remaining whiskers induces altered functional activation patterns that are shown to be related to plasticity and learning processes (Albieri et al., 2015; Alonso Bde et al., 2008; de Celis Alonso et al., 2012; Mirabella et al., 2001; Sellien and Ebner, 2007). Consequently this sensory stimulation paradigm is expected to modulate resting state connectivity, yet it should still be subtle enough to pose a challenge for the analytical methods under evaluation.

One major weakness in investigating resting state fMRI is the heterogeneity of analytical approaches, especially given that the method used to analyze RS data strongly influences the results (Cao et al., 2014; Long et al., 2008; Ma et al., 2007; Rosazza et al., 2012). Therefore, reproducing and comparing findings across different laboratories and studies is difficult. Classical methods to analyze RS data include: regional homogeneity analysis (Zang et al., 2004), cluster analysis (Lee et al., 2012; van den Heuvel et al., 2008a), seed based correlation analysis (SCA) (Greicius et al., 2003) and independent component analysis (ICA) (Calhoun et al., 2001). More recently, graph-theoretical approaches are used, which translate RS data into networks consisting of nodes and edges by cross correlating regional time-courses (Smith et al., 2013). Definition of regions used in the time-course analysis ranges from voxels (Fransson et al., 2011; van den Heuvel et al., 2008b) to only a few anatomical regions (13 nodes) (Fair et al., 2008). The large variance of parcellation resolutions further complicates the ability to compare graph-theoretical findings (Fornito et al., 2010; Wig et al., 2011). Differences in parcellation size influence not only the signal to noise ratio of the mean time-course, but larger parcellation regions are also more likely to integrate more functionally distinct parts.

As illustrated in Fig. 1, RS analysis methods can be classified by their empirical regime (data or hypothesis driven procedures) and their integrated coherence (Long et al., 2008), which is the spatial similarity of all correlating time-courses. Hypothesis driven methods usually involve the definitions of regions of interest (ROIs). These ROIs can be defined either anatomically (e.g. atlas regions) or functionally (e.g. the activated regions of event or stimulus related fMRI). Several reviews provide a comprehensive overview of all methods including the discussion of their advantages and disadvantages (Lee et al., 2013; Margulies et al., 2010).

**Fig. 1:**
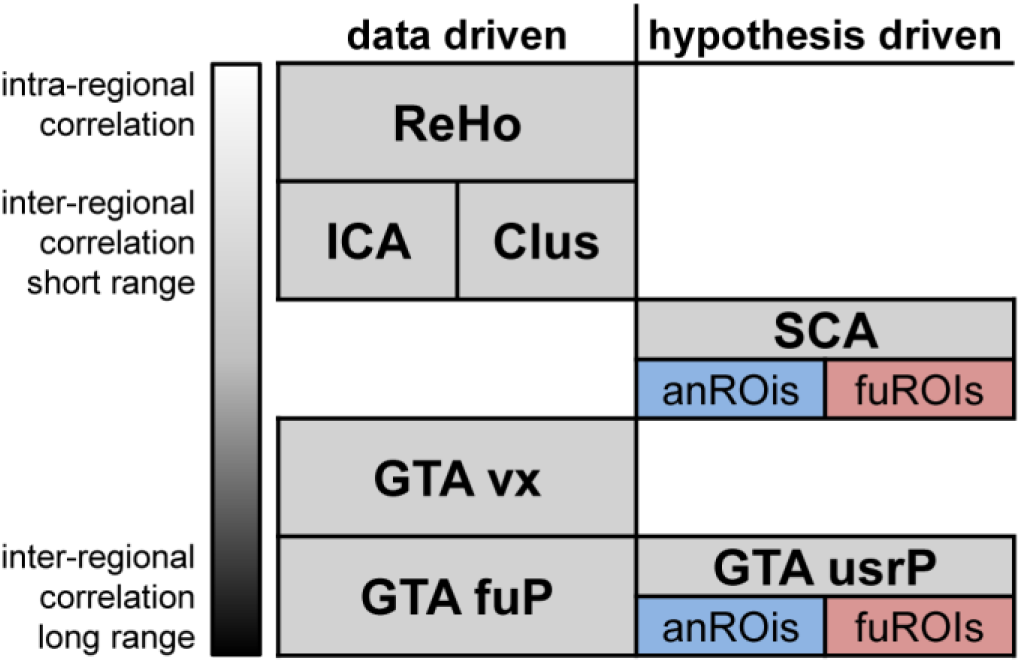
Prominent categories of resting-state fMRI data analysis classified according to their empirical regime and integrated coherence. Data driven methods ranging from intra-regional to long range inter-regional correlations are ReHo: regional homogeneity, Clus: Cluster analysis, ICA: Independent component analysis, GTA vx: voxelwise graph-theoretical analysis, GTA fuP: Graph theoretical analysis based on functional parcellation (Eickhoff et al., 2015; Fan et al., 2016). Hypothesis driven methods are SCA: seed correlation analysis, GTA usrP: graph-theoretical analysis with user defined parcellation. For the last two methods, regions can be defined relying either on anatomical (anROIs) or functional (fuROIs) properties.

In this study we introduce a new method that combines classical seed correlation analysis with graph-theory. We chose SCA as fundamental part of the new approach because seed regions can be small and equal in size providing comparable signal to noise ratios and minimizing the effects of averaging functionally different voxels. Additionally, the correlating voxels in target brain regions are not predefined but determined data driven. That means the method finds for each subject the optimal correlation within the target region. This effect should diminish the variability and enhance the reliability of functional connectivity weights between two regions in brain networks across subjects. We compared this new method to two other graph-theoretical methods that rely on the same parcellation scheme as well as to ICA. Since ICA was the first method used to characterize distinct resting state networks in a data driven fashion, we used it to evaluate the communities obtained with the graph-theoretical methods. We evaluated the efficacy of these methods to characterize short term modulation by comparing RS before and after unilateral whisker stimulation in rats. For this purpose a version of the network based statistics (NBS) (Zalesky et al., 2010) was adapted to cope with pairwise statistics.

## 2. Materials and methods

### 2.1 Animal preparation

Permission for animal experiments was obtained from the ethics committee of the government of Mittelfranken (Ansbach, Germany, 621-2531.31-30/00). fMRI experiments were performed on rats (male Sprague Dawley, 350 to 450 g, Janvier, France, n=25). The animals were initially anesthetized with 5% isoflurane for 7 min in a 1:1 mixture of oxygen and pressured air. Immediately after, rat whiskers were trimmed on both sides of the snout with exception of the ones in the C row. Rats were then mounted on a Plexiglas cradle with incorporated mouth mask and with a tooth biting bar where the rat's head was fixed without the need of ear screws [see de Celis Alonso et al. (2012) for details on experimental setup]. Lateral openings in the mask allowed the whiskers to move freely.

Afterwards the animals were placed in the scanner and the isoflurane concentration was reduced to ≈1.2%. In order to control the depth of anesthesia the isoflurane concentration was adjusted during the experiment to maintain a respiration rate of approximately 65/min (~ 38 mmHg+- 10% pCO2). A stable spontaneous respiration rate leads to stable transcutaneous pCO2 during the fMRI measurement (Ramos-Cabrer et al., 2005). Body temperature was maintained at 37°C by a circulating water bath, and physiological parameters (respiration, temperature) were monitored [for more details see Hess et al. (2011); Hess et al. (2007)].

### 2.2. Experimental protocol

Each animal underwent one fMRI-session starting with a resting state (RS) measurement followed by a stimulation experiment (either sham or whisker stimulation) and a second RS measurement and a final anatomical scan. Animals were separated into two groups, one with stimulation of the remaining whiskers after trimming between the two RS measurement (experimental group, n=13) and one control group prepared and mounted in the same way as the experimental group but without whisker trimming or stimulation (control group, n=12) (Fig. 2).

**Fig. 2:**
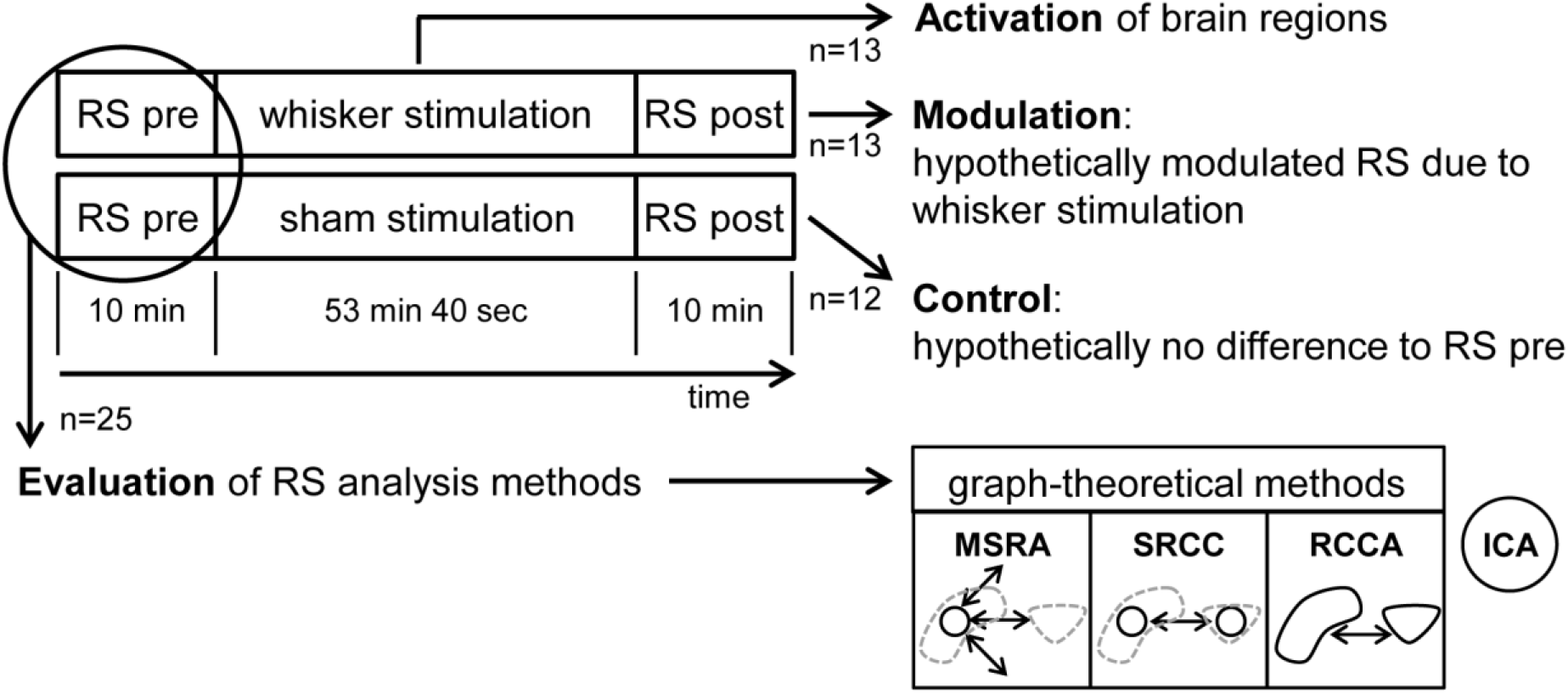
Experimental design including resting state (RS) analysis methods. fMRI images were taken of the animals during three experimental periods per session: RS pre, the stimulation period and RS post. 13 animals were whisker stimulated and 12 were in the control group. The pooled animals of both RS pre periods (n=25) were used to evaluate the RS analysis methods: MSRA (Multiple Seed Region Analysis): time-courses extracted from small seed regions in the center of mass of anatomical brain structures, voxelwise correlated. SRCC (Seed Region Cross Correlation): time-courses extracted from seed regions (see above), cross-correlated. RCCA (Regional Cross Correlation Analysis): time-courses extracted from anatomical brain structures, cross-correlated. ICA: Independent Component Analysis. For detailed description of graph theoretical RS analysis methods see 2.9.

### 2.3 Whisker stimulation

Whiskers were stimulated with an air driven device integrated into the holding cradle. An inverted comb situated 2 cm apart from the left side of the snout was used for monolateral stimulations at a frequency of 6 Hz with an amplitude of 10 mm. All the remaining whiskers from the left side of the snout were stimulated. Space between comb teeth was enough to leave some flexibility for the whiskers to slide in and out, but not to get free, thereby avoiding painful pulling stimuli. Combs were driven from an external console connected to the scanner running a custom programmed user interface developed in LabView (Labview, National Instruments, Austin, TX, USA).

### 2.4 MRI acquisition

MRI experiments were performed on a 4.7 T/40 cm horizontal bore magnet BioSpec (BRUKER, Ettlingen, Germany). A whole-body birdcage resonator enabled homogenous excitation operating with an actively shielded high-power gradient system (200 mT/m) and a low-noise, actively RF-decoupled 2x2 phased array head coil (Bruker Biospin, Ettlingen, Germany) was used to acquire brain images. This configuration enables image acquisition with a high temporal signal-to-noise ratio (tSNR) of approximately 60 (Kalthoff et al., 2011). RS data (collected during both pre and post stimulation period) consisted of 300 brain volumes in each 10 min (cf. Fig. 2) acquired with a T2*-weighted single-shot gradient echo-based Echo Planar Imaging sequence (GE-EPI) covering 22 axial slices of the brain in 2 seconds (total time 10 minutes, TE_ef_ =25.03 ms, TR= 2000 ms, in-plane resolution 0.391×0.391 mm, matrix 64 ×64, FOV 25x25 mm, slice thickness 1 mm). Slice 14 was positioned at bregma -3.48 mm according to Paxinos rat brain atlas (Paxinos and Watson, 2007). As a positioning reference we used the smallest distance between the posterior tip of the corpus striatum and the anterior tip of the hippocampus on the horizontal anatomical reference image.

Both sham and whisker stimulation driven fMRI data (between the two RS measurements) was acquired using the same imaging sequence (GE-EPI). A total of 1602 brain volumes were scanned during this experimental period (cf. Fig. 2) with the following scanning parameters: TE_ef_=25.03 ms, TR=2000 ms, in-plane resolution 0.391×0.391 mm, matrix 64×64, FOV of 25x25 mm, 1 mm slice thickness. The stimulation protocol included 100 stimulations (6 Hz) of the vibrissae with duration of 8 sec (4 volumes) and intermediate rests of 24 sec (12 volumes). First stimulation sequence started after 8 sec. Finally, 22 corresponding anatomical T2 reference images (RARE, RF = 8, TE_ef_ =11.7 ms, TR = 3000 ms, NEX=5, in-plane resolution 0.097×0.097 mm, matrix 256×256, field of view 25×25 mm, slice thickness 1 mm) were acquired at identical positions.

### 2.5 Analysis of BOLD activation due to whisker stimulation

In this study we focused on alterations of RS networks as a result of whisker stimulation. Additionally, the brain structures activated by the whisker stimulation were compared to those involved in resting state connectivity (cf. Fig 2).

BOLD activation induced by whisker stimulation was analyzed using standard procedures described by de Celis Alonso et al. (2012). After appropriate preprocessing including inter-slice-time correction, motion correction, and spatial and temporal smoothing, a general linear model (GLM) analysis was performed (for detailed information see Supplementary Methods). The significantly (FDR, q=0.05) activated voxels in each defined brain structure were determined for each animal. Finally, the number of activated voxels per brain structure was averaged across animals and expressed as percent of total area size of that brain structure.

### 2.6 Preprocessing of RS data

We used Brainvoyager QX 2.8 (Brain Innovation B.V. Maastricht, The Netherlands) for initial inter-slice time and motion correction of RS data. Inter-slice time correction was calculated in ascending interleaved scan order with cubic spline interpolation, and motion correction was computed using rigid registration and resampling with trilinear/sinc interpolation. If not stated otherwise, all subsequent analyses were performed with MagnAn (BioCom, Uttenreuth), an IDL application (Exelis Visual Information Solutions Inc., a subsidiary of Harris Corporation, Melbourne, FL, USA) designed for complex image processing and analysis with emphasis on MR imaging.

After 3D gaussian smoothing (FWHM 1.17 mm), time series were low pass filtered with a frequency of 0.1 Hz and the global signal mean removed by linear regression.

The high resolution anatomical reference images of all 25 animals were co-registered to one selected reference subject using an affine linear transformation algorithm with 6 degrees of freedom and averaged. This mean reference was interpolated to the isotropic voxel size of 0.1x0.1x0.1mm^3^ (to fit requirements of ICA software) and served as the anatomical template. Afterwards, the motion corrected functional RS data of each animal were manually skull-stripped, registered to the 3D anatomical template and linearly interpolated to the same isotropic voxel size.

### 2.7 ICA

Group ICA was performed by using GIFT software (GIFT v1.3g; icatb.sourceforge.net). Briefly, the whole dataset consisting of the concatenated preprocessed time series of all animals was reduced by means of PCA and subsequently decomposed in a predefined set of group-independent components using the Infomax algorithm. Finally, the corresponding components for each animal were calculated by back reconstruction (Calhoun et al., 2001). We calculated group ICAs with 15, 20 and 30 components and evaluated the resulting components by visual inspection. Decomposition into 20 independent components revealed the best correspondence to previous published rodent ICA components (Becerra et al., 2011; Jonckers et al., 2011). Thus, we focused on ICA with 20 components.

### 2.8 Functional connectivity

The analysis of functional connectivity was performed separately on each animal in its native space using the first volume of each RS time series as individual anatomical reference. Regions of interests corresponding to distinct brain structures must be defined prior to running graph-theoretical analyses for functional connectivity. For this purpose we used an in-house digital 3D rat brain atlas with 179 brain structures according to (Paxinos and Watson, 2007). This atlas was semi-automatically registered (affine, 6 degrees of freedom) to each dataset.

A seed region for every brain structure was defined automatically as the 4 voxels nearest to the center of mass, resulting in a cluster of 5 voxels. The following restrictions were taken into account: (i) the voxels must be located within the border of the brain structure; (ii) seed voxels must be located within the central plane of the brain structure as the native datasets are highly anisotropic in the z-direction (slice thickness).

Functional connectivity was represented by the correlation of regional fMRI time-courses stored in a correlation matrix. The definition of the voxels used to extract the time-courses strongly influences the resulting correlation matrices. Therefore, we pooled the animals of the first RS measurement (RS pre) of both groups (n=25) to compare three different methods to extract the time-courses for the creation of the correlation matrices (cf. Fig. 2)

### 2.9 Graph-theoretical methods

The method introduced in this study, is a **multi-seed region approach** (MSRA). It relies on multiple seed correlation maps: the mean time-course of each seed region was correlated with the time-course of every voxel in the brain resulting in one correlation volume per seed region. Subsequently, significant correlations within that correlation volume were determined using false discovery rate (FDR, q=0.05). For each seed region the significant correlation values were averaged per atlas brain structure, resulting in a 179x179 asymmetric correlation matrix (Fig. 3). This asymmetry simulates directionality because for each pair of brain structures, the significant correlation between the seed region and data driven target voxels in the other brain structure is used to generate the correlation matrix. In contrast to the user defined seed regions, target voxels vary within regions and across animals. Note that the correlation itself between seed region and each target voxel is not directed. The resulting asymmetric correlation matrix does not reflect direction in causality, therefore it is called “pseudo directed”.

**Fig. 3:**
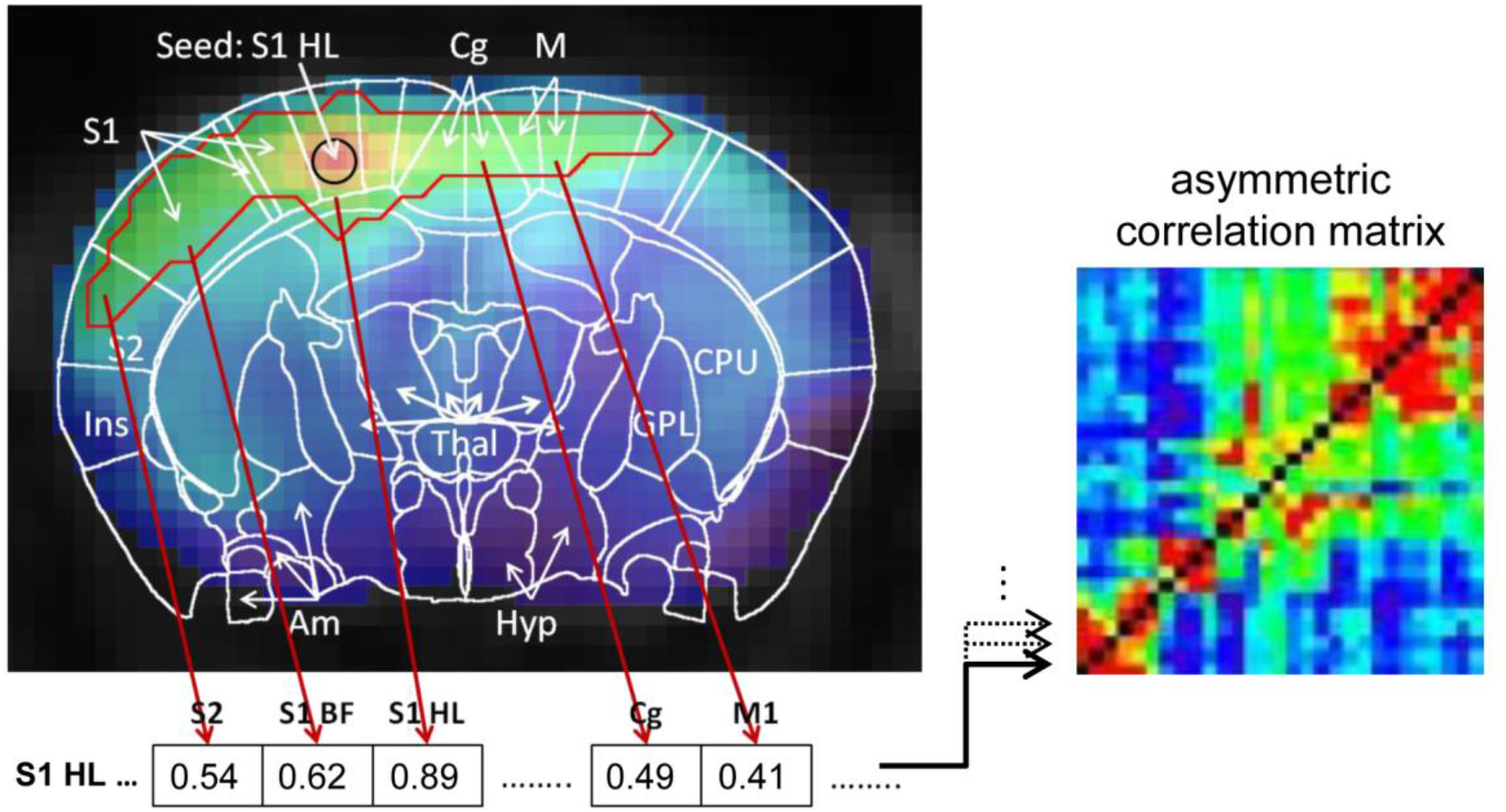
Schematic demonstration of the MSRA-correlation matrix computation using the example of the seed region in the center of mask of the hind limb field of the primary somatosensory cortex (S1HL). Significant correlations of the seed’s correlation map were determined using false discovery rate (FDR, q=0.05, red outline). For each brain area covered by the red outline the significant correlation were averaged and stored in a matrix line. The matrix line values for all other brain areas are set to 0. This procedure was repeated for every seed region resulting in an asymmetric correlation matrix where every line represents a seed region and its average significant correlation with every other brain area.

The second method, **a seed region cross-correlation** (SRCC) analysis, simply cross correlates the mean time-courses of all seed regions (identical to those used in MSRA) to each other, resulting in one symmetric correlation matrix per subject.

The third method, a **regional cross-correlation approach** (RCCA), correlates the mean time-courses of the whole brain structure to each of the other brain structures. However, it is biased by the highly varying numbers of voxels used to create the average time-course per brain structure. This can lead to false correlations, especially in inhomogeneous regions. To weaken this effect, we used the first principal component of each region instead of the simple mean to calculate a cross correlation matrix for each subject (Pawela et al., 2008; Zhong et al., 2009). Briefly, principle component analysis (PCA) is a multivariate technique that replaces the measured variables (here the time-courses of the voxels in the region) by a new set of uncorrelated variables (principle components), arranged in order of decreasing variance. The mean variance that could be explained by the first principal component for all animals and structures was 60 %±16%.

In accordance with the MSRA approach only significant correlations in the SRCC and RCCA correlation matrices (FDR, q=0.05) were counted as connections. In case of the MSRA method the FDR correction was applied to the whole brain, in case of SRCC and RCCA to the correlation matrix. The resulting FDR thresholds are comparable (mean r of all animals ± standard deviation: MSRA 0.1754 ± 0.004, SRCC 0.1742 ± 0.003, RPCC 0.1723 ± 0.002). Non-significant connections were set to 0. Finally, the resulting Pearson’s r correlation values were transformed into Fisher’s z-values to provide normal distributions for subsequent statistical analysis.

### 2.10 Network graphs

To obtain representative group correlation maps, we averaged the individual correlation matrices of all subjects for each graph-theoretical approach. These average correlation matrices were transformed into network graphs consisting of vertices (or nodes) and edges. Vertices represent the brain regions, whereas edges between pairs of brain regions indicate their functional connectivity. The topology of a network graph is strongly dependent on the number of represented connections.

The connections configuring a graph with a given sparsity were defined by thresholding the associated correlation matrix with a fixed threshold or a fixed number of connections. The former procedure accounts for the overall strength of the matrix and results in different numbers of connections per graph (i.e. different sparsity), the latter leads to different threshold values per matrix but equal sparsity and therefore comparable graph-topology. Here it should be considered, that in undirected graphs (RCCA and SRCC) each connection accounts for two directed connections (MSRA).

Network communities were detected using a heuristic method that is based on modularity optimization proposed by (Blondel et al., 2008) and implemented in NWB [NWB Team (2006). Network Workbench Tool. Indiana University, Northeastern University, and University of Michigan, http://nwb.slis.indiana.edu]. The nodes within these communities are more strongly connected to each other than to nodes outside the community. Since we intended to compare the network communities resulting from the different graph-theoretical approaches with the 20 ICA components of the same dataset we limited the number of connections to 20 communities per approach (MSRA: 600 directed connections, RCCA: 300 undirected connections and SRCC: 385 undirected connections). The networks were visualized in Amira 5.4 (Visage Imaging) using a force-based algorithm (Kamada and Kawai, 1989).

The sensitivity of the approaches to robustly detect changes in RS networks was evaluated by investigating the short term modulation of RS network graphs during whisker stimulation. For this purpose the correlation matrices calculated from the first and second RS measurement for the two groups (experimental group with and control group without whisker stimulation, n=13 and n=12, respectively), were averaged separately resulting in 4 averaged correlation matrices per graph-theoretical approach. From these matrices network graphs were created using the 1790 strongest directed connections for MSRA and 895 undirected connections for RCCA and SRCC. This resulted in 20 directed or 10 undirected connections per node. These conditions allow graphs that are fully connected, yet sparse enough to show a topological network organization clearly distinct from that of a random network.

### 2.11 ICA co-activation index

To evaluate the correspondence between ICA and each graph-theoretical analysis, we calculated an “ICA co-activation index” as proposed by (Rosazza et al., 2012). The z-scores (resulting from the GIFT ICA analysis) of each brain region were averaged and multiplied for each pair of brain regions resulting in a 179x179 product matrix. This procedure was performed for each ICA component separately and the product matrices of all ICA components were summed. Adding a power factor k to the procedure described above emphasizes either the relative weight of intense (k>1) or less intense (k<1) co-activation [for details see (Rosazza et al., 2012)]. For the calculation of the ICA co-activation matrix we applied the same methodological constraints used to create the graph-theoretical correlation matrices: (1) only positive z-scores were considered and negatives set to 0 and (2) for each subject a separate ICA co-activation matrix was calculated and subsequently averaged to create a group co-activation index.

### 2.12 Similarity

We used two different approaches to measure the similarity for the three graph-theoretical methods and their correspondence to ICA: (i) the overall linear Pearson’s correlation coefficient r for the comparison of the matrices and (ii) the Jaccard index for the comparison of brain structure lists of the graph-theoretical communities and the list of brain structures comprised in the ICA components.

The Jaccard index measures similarity between finite sample sets. It is defined as the size of the intersection divided by the size of the union of the sample sets. Each similarity measure was calculated for the graph-theoretical methods to each other (the overall r coefficient on single animals, the jaccard index on group level) and for each graph-theoretical method with ICA (both similarity measures on group level). For the latter comparison, the average ICA co-activation index matrices for power levels 0.5, 1 and 2 were used to calculate the overall r coefficient and the lists of brain structures comprising the 5 most stable group ICA components (binarized to z > 0.3) were used to calculate the Jaccard index. For the graph-theoretical communities, the brain structure lists used to calculate the Jaccard index vary with the number of graph connections. Therefore, the comparison of graph-theoretical methods to each other was performed on graphs with 600 to 6000 connections. For the comparison to ICA, we used graphs with connection numbers that resulted in 20 communities to match the number of ICA components (see section 2.8). For statistical analysis of overall r coefficients paired t-test (among graph theoretical methods) and ANOVA (group level graph theoretical methods with ICA) were performed with α < 0.05.

### 2.13 Reproducibility

Reproducibility of graph-theoretical approaches and ICA co-activation matrix was evaluated using the variance matrices of all analysis methods. For each pair of brain areas, the variance of correlation z-value was calculated and normalized to the squared mean z-correlation value. The median of all normalized correlation variances demonstrates the reproducibility of each analysis approach. The confidence-interval of each median was calculated by bootstrapping. Significant effect of methods was tested using the Kruskal-Wallis test. Additionally, mean-subtracted variance matrices were calculated, which show the variability of variance within all pairs of brain areas.

### 2.14 Paired network based statistics (pNBS)

To statistically evaluate the short term modulation of RS network graphs during whisker stimulation, we implemented a modified version of the network-based statistics (NBS) introduced by (Zalesky et al., 2010). The NBS relies on the assumption that group differences in single connections are more likely to be false positive than differences in larger connected components. To each connected component of group differences, a p-value controlled for the family-wise error can be ascribed using permutation testing. For details see Zalesky et al. (2010).

In experimental designs with paired groups, permutations cannot be used. To overcome this limitation we included an additional control experiment with identical experimental parameters with the exception of stimulation between the two RS measurements (cf. Fig.2). Our pNBS approach uses the fisher’s z-transformed correlation matrices of all four RS scans per animal (2 for experimental and 2 for control group). For each animal, the differences of correlation values per connection between the two RS correlation matrices of each session (control and experimental) were calculated. These difference matrices were used to calculate the pairwise T-test statistics and for the permutation testing.

The following steps were used to determine modulated connected components (1) Paired t-statistics was computed for both control and experimental group. (2) The 99% quantile of the p-values of the paired t-statistics of the control group was determined to identify a set of supra-threshold links corresponding to 1 % hypothetical false positive connections (we hypothesize there are no stimulus dependent connectivity modulations in the control experiment). (3) The same threshold was applied to the paired t-statistics p-values of the experimental group, all connected components above this threshold equal or smaller to the largest component of the control group were eliminated. The size *k* of the remaining set of connections was stored. In contrast to the traditional NBS these connections might comprise more than one connected component. Finally, a p-value controlling for the family-wise-error was ascribed to the remaining component based on its size using permutation testing.

In each of *M* permutations, the difference matrices were randomly assigned to either control or experimental group, and the statistics of interest described in steps (1) to (3) was recalculated. The size *k’* of the set of supra-threshold links derived from each permutation was determined and stored. The p-value of an observed set of connections is estimated by finding the total number of permutations with *k’* > k normalized by *M*.

## 3. Results

### 3.1 Positioning of seed regions

One critical step during seed region based analysis of RS data is the reproducibility of positioning the seed regions across subjects. The MSRA approach includes an automatic positioning step for all seed regions (see section 2.9). Fig. 4 demonstrates the reproducibility of this procedure. Although all animals were individually analyzed (including atlas registration and automatic determination of seed regions), the positions of the seed regions after affine registration of all animals remained consistent. The Euclidian distance to the mean centroid per structure was on average 1.4 pixel (0.55 mm) in x-y plane and 0.2 slices (0.2 mm) in z direction.

**Fig. 4:**
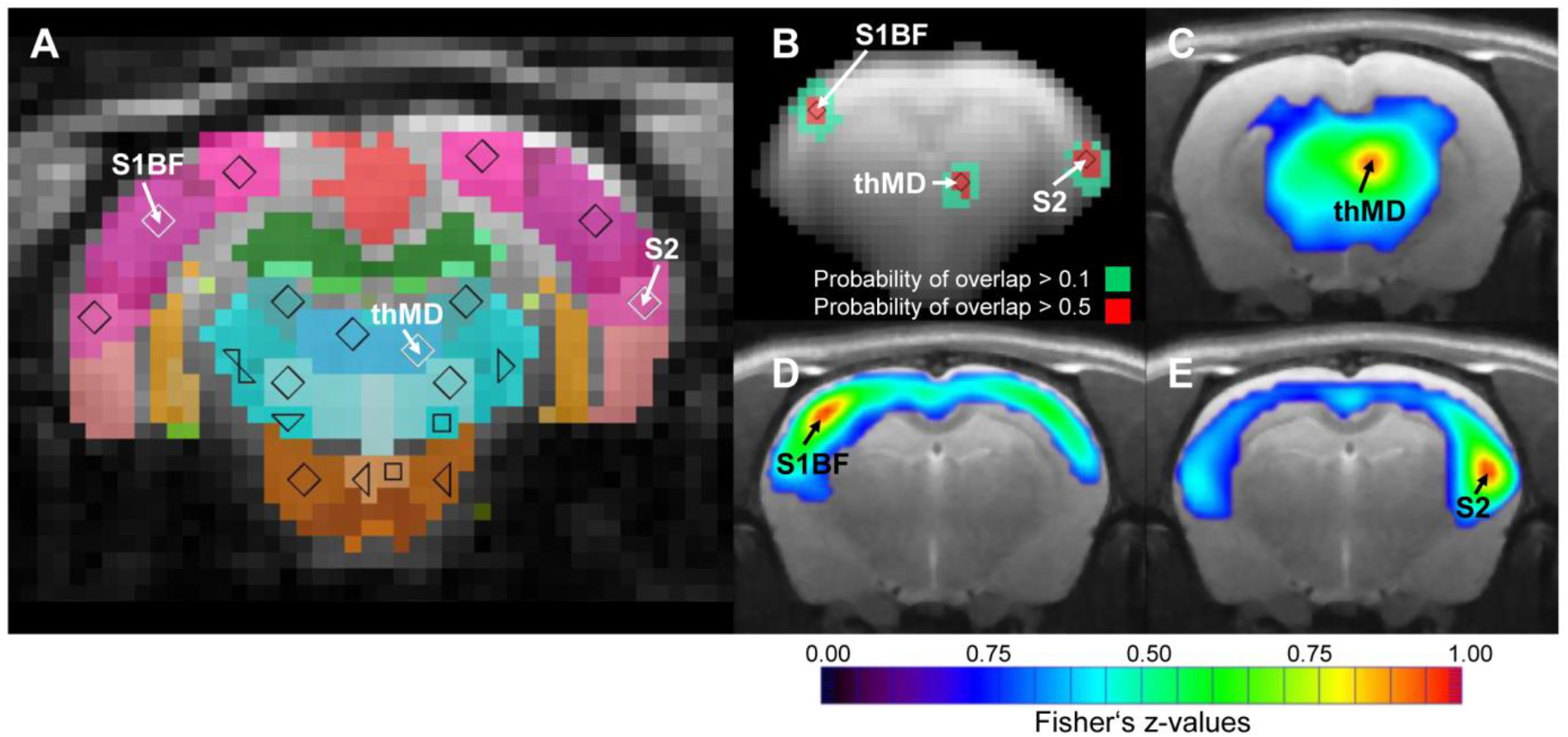
Consistency of automatically placed seed regions of the MSRA approach. A) Seed regions in the center of mass of individually matched digital atlas regions of one animal. B) Overlap of exemplary binarized seed region voxels (n=25). C-E) Up-sampled high resolution correlation maps of exemplary seed regions. S1BF: barrel field of primary somatosensory cortex, S2: secondary somatosensory cortex, thMD: mediodorsal thalamus.

### 3.2 Comparison of methods: Correlation matrices

The correlation matrix of the MSRA approach differs from both SRCC and RCCA, which are comparable to each other (upper triangles in Fig. 5A). This effect is associated with the different distribution of z-values within the correlation matrix. SRCC and RCCA matrices show left shifted distributions with their maxima below z=0.01, which is below the significance level of the FDR (average over all animals: z=0.176 and z=0.174, respectively). However, the histogram of the MSRA approach has a maximum around z=0.25 (average FDR z=0.177) (Fig. 5B). The number of connections is determined by thresholding the correlation matrix with a given z-value. For lower z thresholds, the numbers of connections are similar between SRCC and RCCA, but the MSRA approach has considerably more connections. For threshold values higher than 0.4 this effect is reversed resulting in slightly lower number of connections in MSRA compared to SRCC and RCCA (insert in Fig. 5C).

**Fig. 5:**
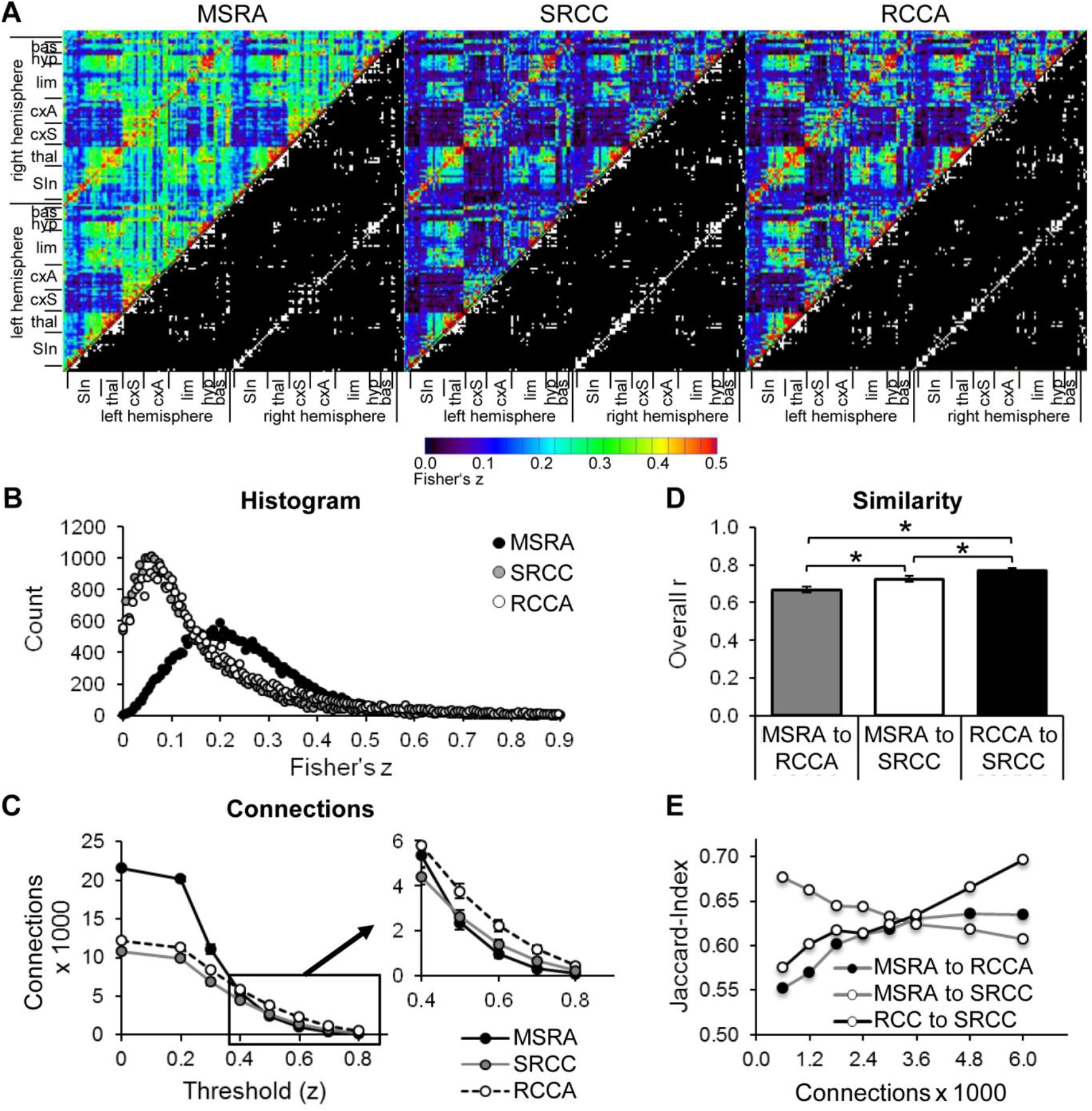
Comparison of graph-theoretical network approaches. A) Correlation matrices including top 1790 binarized directed connections (lower right triangle). B) Histograms of Fisher’s z-correlation values. C) Number of directed connections as a function of the threshold used for binarization (mean +− SEM, n=25). Pairwise similarity of graph-theoretical approaches: D) overall correlation r of correlation matrices (mean +− SEM, n=25, * p<0.05, paired t-test) (E) and Jaccard index as a function of the number of directed connections of binarized average matrices. bas: basal ganglia, cxA: association cortex, cxS: sensory cortex, hyp: hypothalamus, lim: limbic system including hippocampus and amygdala, SIn: sensory input, thal: thalamus.

The similarity of the correlation matrices of the different approaches was pairwise determined by the overall r-coefficient of all weighted connections (Fig. 5D) and by the jaccard index of binarized matrices (Fig. 5E). The overall r reveals highest similarity between RCCA and SRCC. As expected, the overall similarity between MSRA and SRCC is higher than between MSRA and RCCA networks because the MSRA and the SRCC approach rely on the same seed regions (Fig. 5D).

The jaccard index of RCCA and SRCC and RCCA and MSRA networks increases with the number of connections. In contrast, the similarity between MSRA and SRCC networks decreases with the number of connections (Fig. 5E).

### 3.3 Network communities

Of the 20 communities, the number containing at least four nodes is equal for MSRA and SRCC networks (12) but differs for RCCA (15). Five of these RCCA-communities are very small (consisting of only four nodes) indicating slightly higher network segregation. In general, the communities contain nodes that represent anatomically and/or functionally associated brain structures (Fig. 6, Suppl. Fig. S1).

**Fig. 6:**
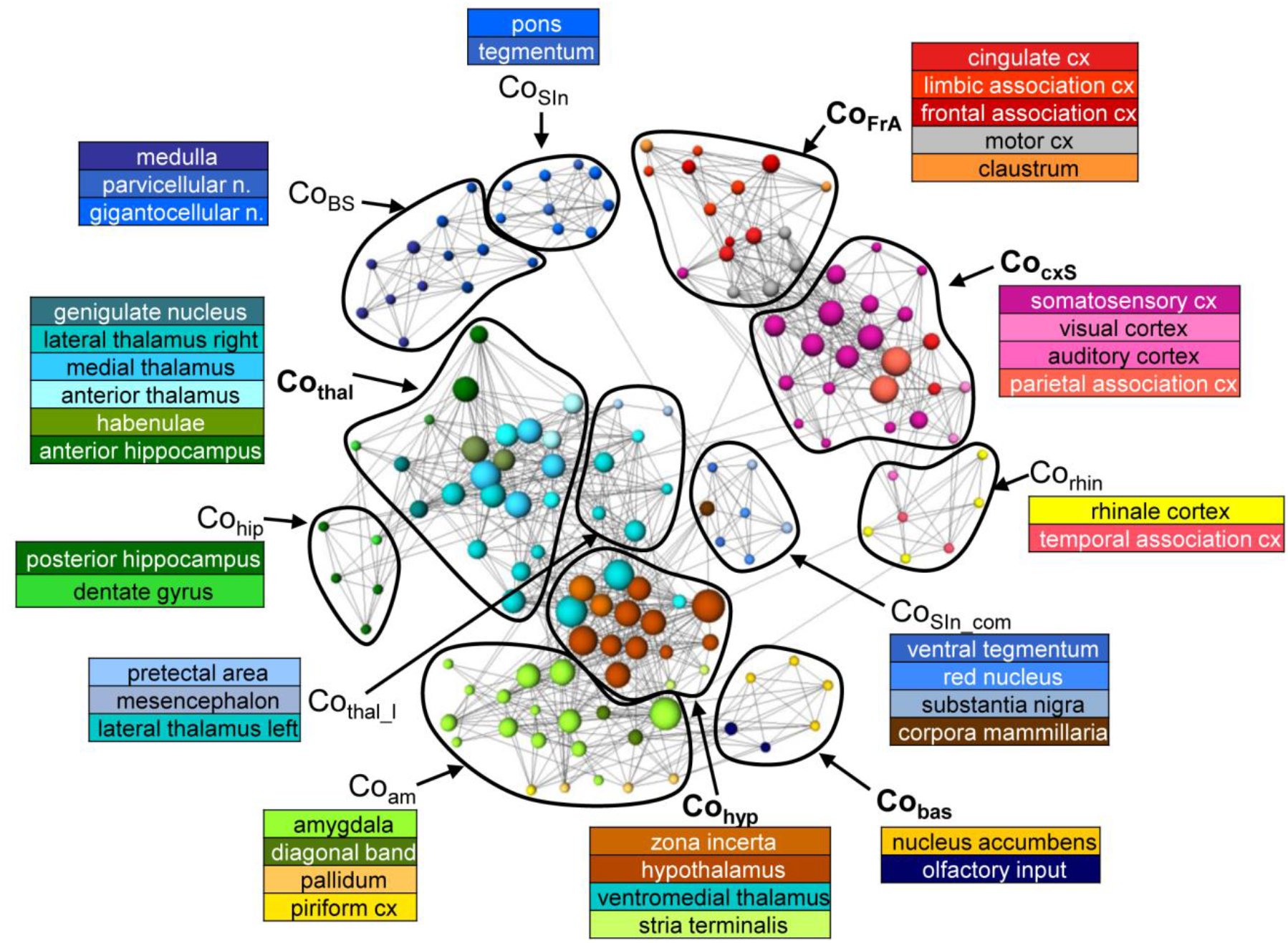
Communities of the average RS network (n=25) from the MSRA approach. Node positions are determined using a forced based algorithm. For better visualization the underlying network comprises 1790 strongest connections and only communities that contain at least 4 nodes are shown.

The MSRA network consists of the following communities (Fig. 6): Brainstem including medulla with solitary tract and parvicellular and gigantocellular nuclei (Co_BS_); sensory input including pons and tegmentum (Co_SIn_); structures of the sensory input (ventral tegmental area, red nucleus, substantia nigra) connected to the corpora mamillaria (Co_SIn-com_); thalamic structures associated with anterior hippocampus and septum (Co_thal_) or with mesencephalon and pretectal area (Co_tha1_1_); structures of the sensory cortex (S1, S2, visual and auditory cortex) and the parietal association cortex (Co_cxS_); frontal association areas, cingulum, motor cortex and claustrum (Co_FrA_); ecto- and entorhinal cortex (link to the limbic system) in community with the temporal association area (Corhin); posterior hippocampus and dentate gyrus (Cohip); limbic structures (mainly amygdala), pallidum (basal ganglia) and piriform cortex as link to limbic system (Coam); hypothalamus and zona incerta associated with the ventromedial thalamus and stria terminalis (Cohyp); nucleus accumbens and olfactory nucleus (Co_bas_).

The most striking deviation of the SRCC-network communities with respect to the MSRA is that the hippocampal areas are segregated into left and right hemisphere and are more strongly connected to thalamic structures of the same hemisphere (Suppl. Fig. S1A). The communities of the RCCA network show even more deviations from the MSRA network communities described above (Suppl. Fig. S1B). The most prominent difference is the higher degree of segregation. This mainly includes the cortical sensory structures, (corresponding to the MSRA-Co_cxS_-community), but limbic structures (diagonal band and pallidum, septum and stria terminalis) and the medulla are also affected. On the other hand, the frontal association areas and basal ganglia (corresponding to the communities Co_FrA_ and Co_bas_ in the MSRA-network) are integrated into one large community. Here, the motor cortex is associated with the sensory instead of the frontal association cortex.

In summary, the communities tend to be consistent for all three network analysis methods. However, they each produce some different connections resulting in community variations. The MSRA is more similar to the SRCC at the community level as already shown in Fig. 5D and E for the correlation matrices.

### 3.4 ICA Components

We used Independent Component Analysis (ICA) to compare these three methods to an established, independent method,. The following six most stable non-noise ICA components widely reflect the communities of the graph networks (Fig. 7):

**Fig. 7:**
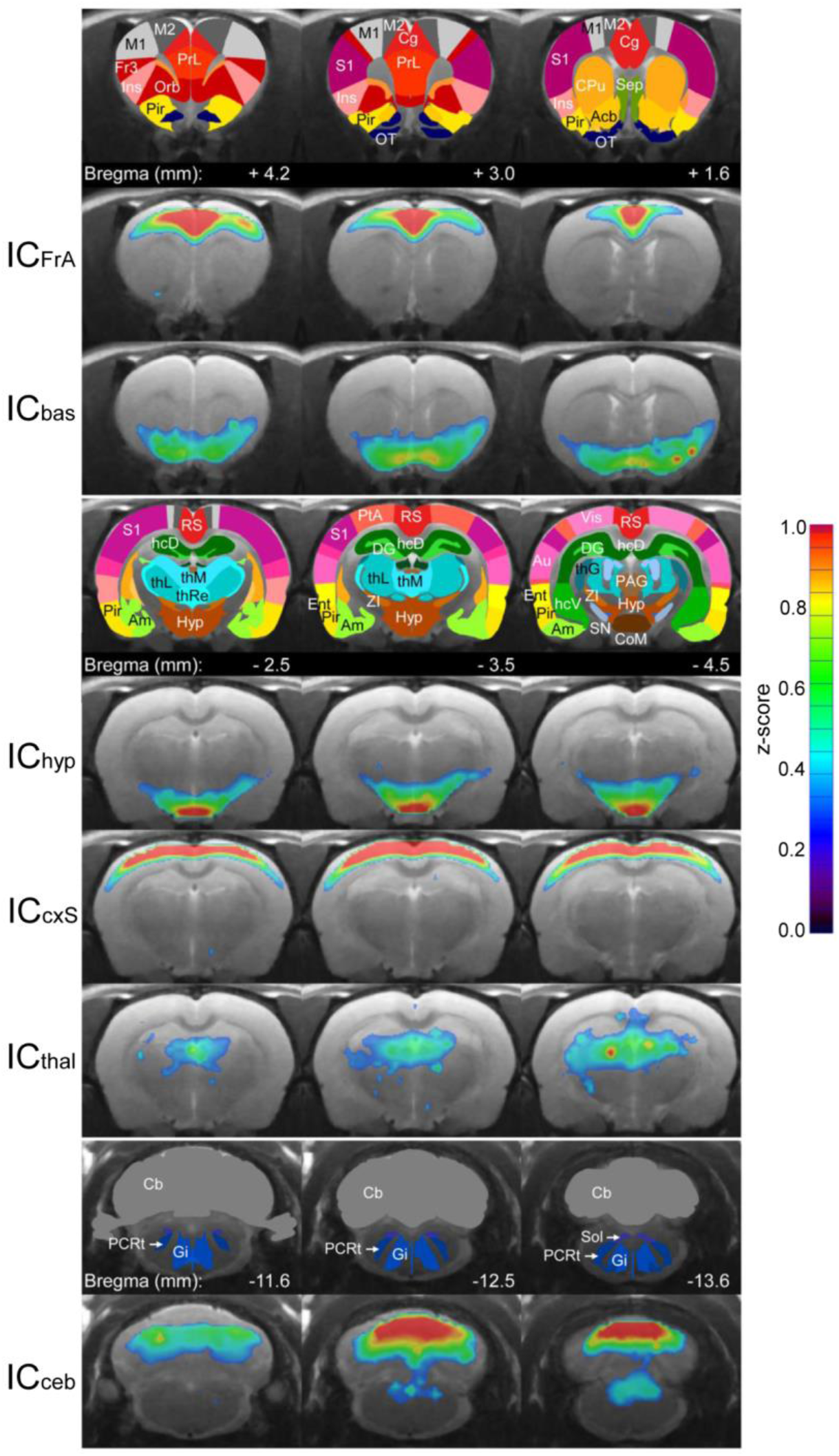
Average (n=25) RS ICA components used for comparison with the graph-theoretical methods. The color-coded ICA z-score maps of these components are overlaid on the anatomical image. Z-scores represent the correlation between each voxel time-course and the mean time-course of the associated component. For each component, three subsequent coronal slices are shown with the corresponding atlas regions in the row above. Acb: nucleus accumbens, Am: amygdala, Au: auditory cortex, Cb: cerebellum, Cg: cingulum, CoM: corpora mammilaria, CPu: caudate putamen, DG: dentate gyrus, Ent: entorhinale cortex, Fr3: frontal cortex area 3, Gi: gigantocellular reticular nucleus, hcD: dorsal hippocampus, hcV: ventral hippocampus, hyp: hypothalamus, Ins: insula, M1: primary motor cortex, M2: secondary motor cortex, Orb: orbitofrontal cortex, OT: olfactory tubercle, PAG: periaqueductal gray, PCRt: parvicellular reticular nucleus, Pir: piriform cortex, PrL: prelimbic cortex, PtA: parietal association cortex, RS: retrosplenial cortex, S1: primary somatosensory cortex, Sep: septum, SN: substantia nigra, Sol: solitary tract, thG: geniculate thalamus, thL: lateral thalamus, thM: medial thalamus, thRe: reunions thalamic nucleus, Vis: visual cortex, ZI: zona incerta.

IC_FrA_: This component includes the prelimbic association cortex, cingulum and motor cortex. It corresponds to the community Co_FrA_ of the MSRA network.
IC_bas_: This component corresponds to the Cobas-Community of the graph network. It includes the olfactory nucleus and the nucleus accumbens as well as the piriform cortex.
IC_hyp_: Comparable to the graph-theoretical Cohyp-Community this IC involves predominantly autonomic regions including hypothalamus and corpora mammillaria.
IC_cxS_: The IC_cxS_ involves sensory cortical structures (S1, S2, auditory and frontal part of visual cortex) and the parietal and retrosplenial cortices. It reflects the CocxS-community.
IC_tha1_: Mainly thalamic structures associated with hippocampal areas and septum are involved in this component.
IC_ceb_: This component is the most caudal one and covers large parts of the cerebellum, which is connected to structures of the brainstem (solitary tract, reticular nucleus). Apart from the cerebellum; which is omitted in graph-theoretical network analysis, this IC is most likely represented by the brainstem community (CoBS). Of note, the ICA components in the rodent brain found in this study align with the ones already published (Becerra et al., 2011; Liang et al., 2011).

### 3.5 Similarity of ICA Components and Network Communities

The similarity of ICA and network-connectivity was determined by the average r of all weighted connections per graph-theoretical approach and the ICA co-activation index matrix at three different power factors (Fig. 8A). Two factor ANOVA revealed a significant effect for the factor “method” (p=0.017) and for the factor “power factor” (p=0.048). The MSRA approach shows the highest overall-r values compared to the other two methods indicating more similarity to ICA components. The maximum similarity occurs at power factor 1. Focusing on either weak (power factor < 1) or strong (power factor > 1) connections causes a decrease in similarity (insert, Fig. 8A) for the MSRA approach but a further increase for power factor > 1.0 for the other two methods.

**Fig. 8:**
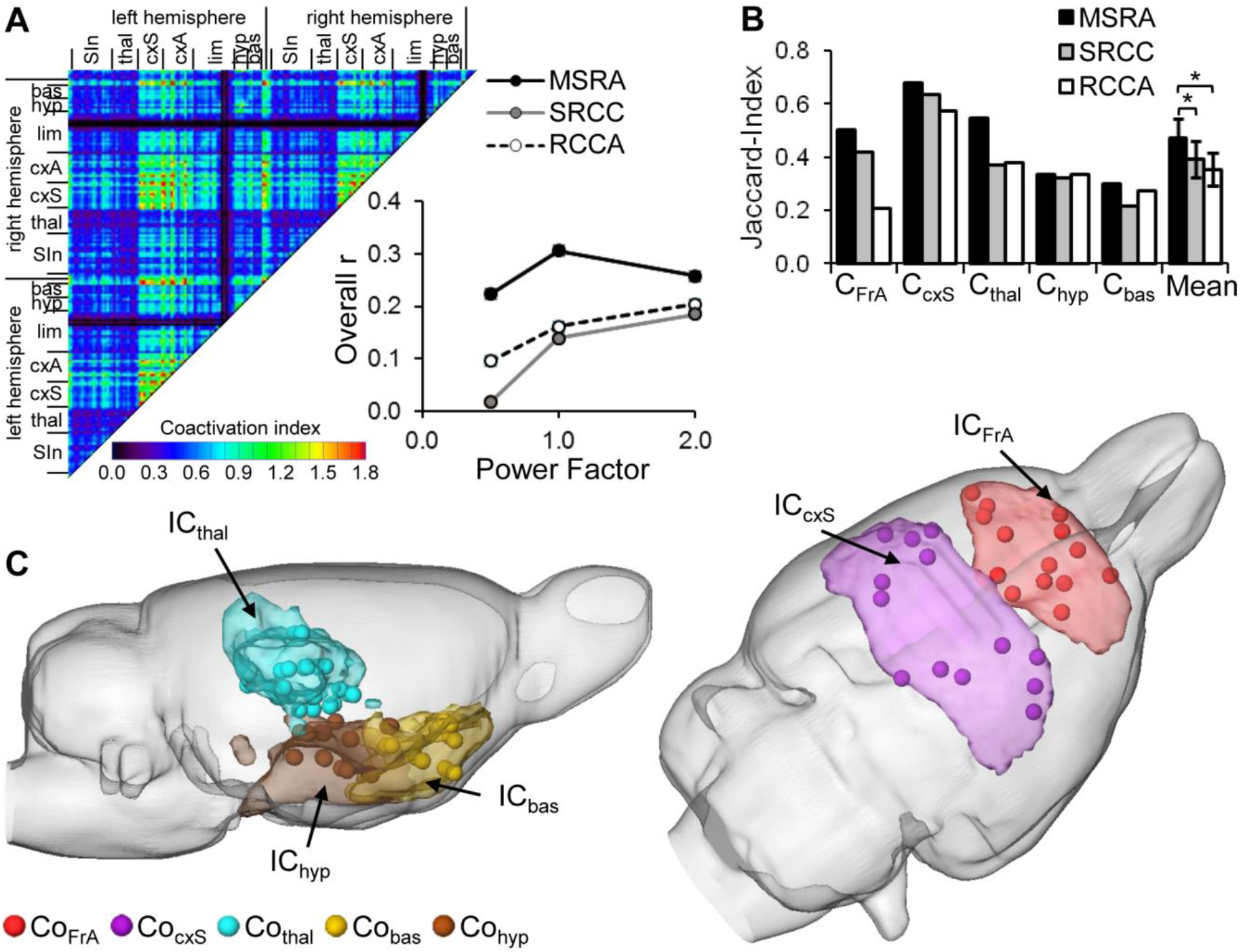
Similarity of RS ICA components (IC) and graph-theoretical network communities (Co). A) ICA coactivation index matrix (power factor 1) and its overall correlation with the average correlation matrices of the three graph-theoretical approaches as a function of power factor. B) Jaccard-Index of brain areas belonging to associated ICA components AND/OR network communities (*p<0.05, paired t-test, n=5 communities). C) Visual overlap of thresholded ICA components (> z=0.03) and centroids of the brain areas composing the associated MSRA network communities.

Additionally, we compared the binarized ICA components with the network communities obtained by the three different graph-theoretical approaches. Because of the missing cerebellum node in the graph-theoretical analysis ICceb and CoBS were not included in the comparison. The overlap was quantified using the Jaccard index (Fig. 8B). For all community/component pairs, the MSRA approach reveals the highest overlap. This effect is manifested in the mean Jaccard index over all five compared community/component pairs. The mean Jaccard index of the MSRA approach is significantly higher than that of the other two methods (p<0.05, paired t-test, cf. Fig. 8B). As illustrated in Fig. 8C, ICA components and the MSRA network communities share a great deal of overlap.

### 3.6 Reproducibility

In order to quantify reproducibility of the three graph-theoretical approaches and ICA, we used the variance matrix of all animal’s correlation (graph theory) and coactivation (ICA) matrices representing the variance for each connection. The median of connection variances differed significantly for all evaluated methods. It was lowest for the MSRA approach, indicating higher reproducibility compared to the SRCC and RCCA methods (Fig.9A).

Interestingly, brain structures at different organizational levels show unequal variances, which are especially obvious for both SRCC and RCCA (Fig 9B). In contrast, the variance distribution of the MSRA approach and the co-activation indices is much more homogenous. For all three graph-theoretical approaches, the highest variances occur in connections between cortex and subcortex, especially in thalamo-cortical connections. The MSRA approach also showed that seed regions placed in cortical regions had more reproducible connections to subcortical structures than vice versa (Fig. 9B).

**Fig. 9:**
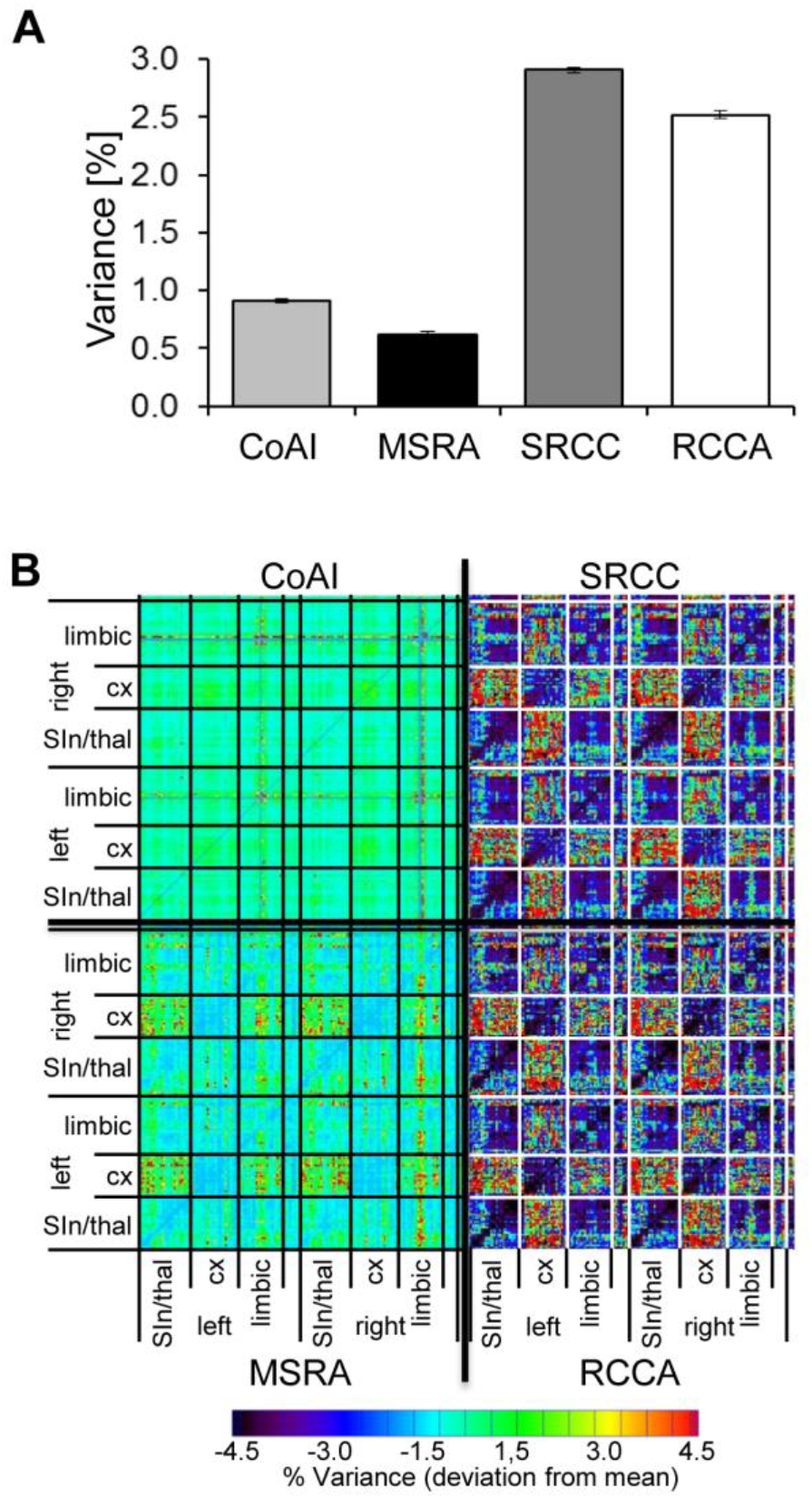
Reproducibility of ICA co-activation index (CoAI) and the three graph-theoretical approaches. A) Medians ± 95% confidence intervals of the variances of all connections demonstrate significantly different reproducibility of analysis methods (p<0.001, Kruskal-Wallis test). B) Mean-subtracted variance matrices demonstrate the variance within all pairs of brain areas.

### 3.7 Short term modulations of RS networks due to whisker stimulation

To determine if the graph approaches are sensitive enough to detect RS modulations, we used them to analyze the RS measurements separated by sensory whisker stimulation in the experimental group compared to the controls. The statistical significance of the induced modulations was determined using pNBS (see section 2.14).

Compared to the control condition (no stimulation between the two RS measurements), whisker stimulation in the experimental group led to significant (p=0.032, corrected using pNBS) alterations in network connectivity (Fig. 10A). Basically, these alterations occurred between brain structures that had been active during whisker stimulation (Suppl. Fig. S2). One exception was the ipsilateral somatosensory barrel field, which did not significantly change connectivity (Suppl. Fig. S2, Fig. 10A). However, the connections of neighboring somatosensory structures in the Co_cxS_-community and the ipsilateral (left) motor cortex were altered, some with increasing (including ipsilateral upper lip field) but most with decreasing connectivity strength. Additionally, subcortical regions were affected, predominantly hypothalamus and ventromedial thalamus (Co_hyp_) with decreasing and amygdala (Co_am_) with increasing connectivity strength (esp. basolateral amygdala). Both hypothalamus and amygdala had weaker connections to the ento- and ectorhinale cortex (Co_rhin_), which represents a link to the limbic system (Fig. 10A). All four structures with the most altered connections (i.e. the biggest nodes in Fig. 10A); the right ventromedial thalamus, right lateral and medial hypothalamus and right basolateral amygdala, were located contralateral to the stimulation side.

**Fig. 10:**
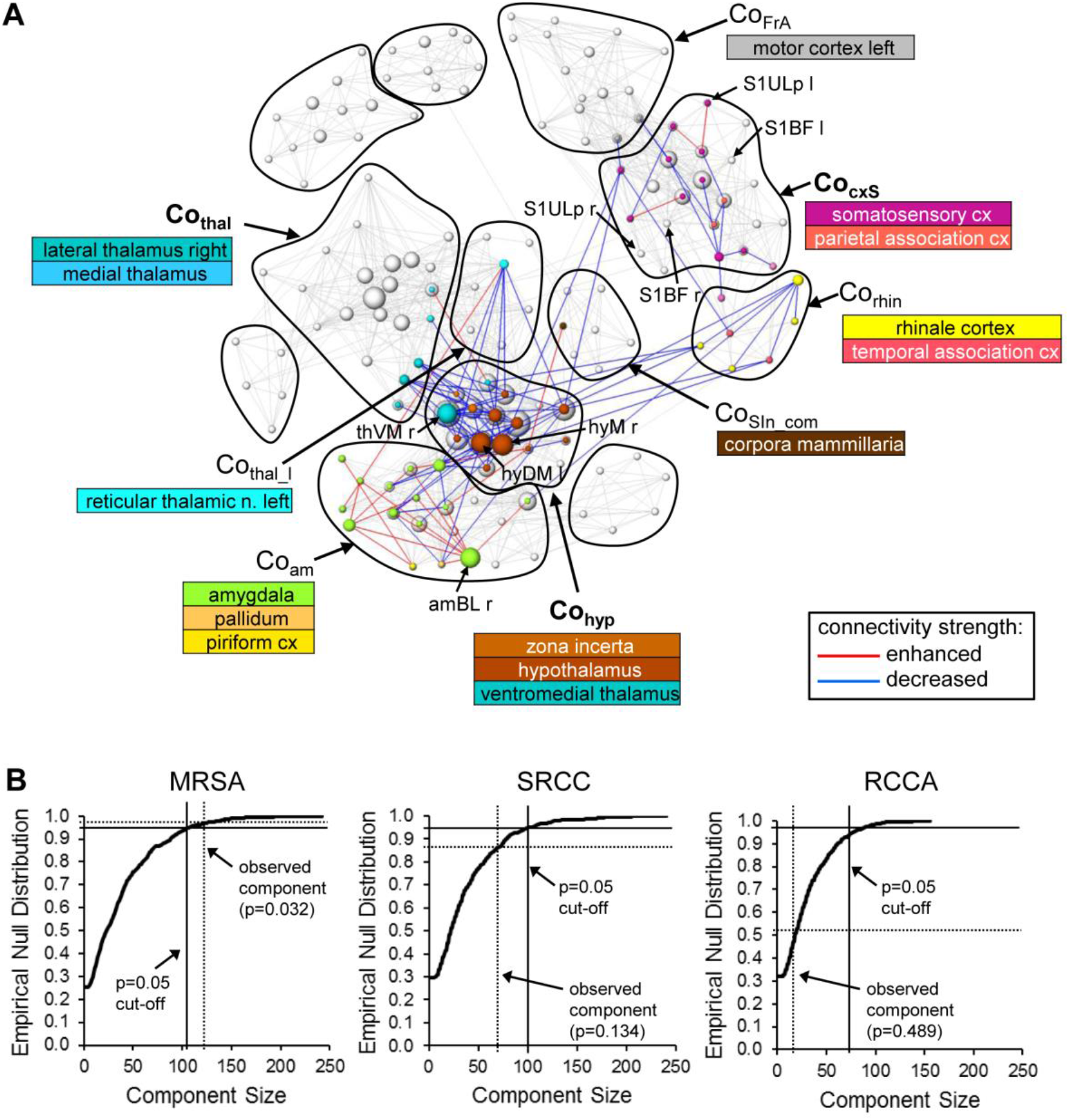
Alterations in RS connectivity due to whisker stimulation. A) Components of connectivity alterations identified by pNBS using the MSRA approach (p=0.032, FWE corrected) overlaid on the RS network before whisker stimulation (grey edges and nodes). The visualization scheme is the same as in Fig. 6. B) Empirically computed null distribution of component size for all three graph-theoretical approaches. Only the observed component of the MSRA was significant (p<0.05) after permutation correction. amBL: basolateral amygdala, hyDM: dorsomedial hypothalamus, hyM: medial hypothalamus, S1BF: primary somatosensory cortex barrel field, S1Ulp: primary somatosensory cortex upper lip field, thVM: ventromedial thalamus, l: left hemisphere r: right hemisphere.

Applying pNBS to graphs obtained with the SRCC and the RCCA method revealed smaller components of altered connectivity compared to the MSRA approach (Suppl. Fig. S3). These components overlap those of the MSRA and thus qualitatively confirm the whisker induced RS modulation effect. However, in contrast to the MSRA approach these two methods did not survive permutation testing. This was demonstrated by the calculated empirical null distribution of each method shown in Fig. 10B.

Additionally, the initial 1% quantile threshold of the paired t-test between the first and second RS of the control experiment was higher for the MSRA approach (p=0.0084) than for the other graph-theoretical methods (0.0032 for both SRCC and RCCA). This effect was in concordance with the lower variance across animals shown in Fig. 9B and confirms the results of the MSRA approach are more reproducible than SRCC and RCCA.

In summary, the MSRA approach, which uses both hypothesis and data driven properties, is superior for graph-theoretical analysis of RS data. Particularly in combination with the proposed pairwise network based statistical evaluation of changes in connectivity strength this approach facilitates the ability to detect and define the slight differences in RS networks induced by a physiological sensory stimulus.

## 4. Discussion

The aim of this study was to introduce a new method for RS analysis that integrates the advantages of traditional ICA and graph-theoretical analysis. The advantages of this new method were illustrated in an investigation of short term modulation of rodent RS connectivity using physiological whisker stimulation.

### 4.1 ICA and seed region analysis

The new MSRA approach relies on multiple seed region correlation analysis steps. It resembles the results of an independent component analysis more than the other two graph-theoretical approaches tested here (SRCC and RCCA), which solely rely on cross correlations of predefined regional time-courses.

Both ICA and classical seed region analysis (SCA) reflect inter-regional features of functional connectivity, whereupon ICA reflects integrated synchronization among networks, SCA measures the inter-regional (long-range) correlation of a specific area with others. Seed based and ICA based RS networks are highly concordant (Barkhof et al., 2014; Van Dijk et al., 2010). For example, the posterior cingulate cortex seed recapitulates the ICA default mode network in human RS analysis (Fox et al., 2005; Long et al., 2008), and seeds within the anterior insula can be used to render the ICA salience network (Seeley et al., 2009).

Differences between ICA and seed based analysis are also reported. (Ma et al., 2007) compared both analysis methods on human fMRI data and on simulated data with artificially added gaussian distributed random noise and a (human) cardiac signal to simulate structured noise. They found that ICA differs from seed based analysis in terms of extent and location of the detected networks. Additionally, ICA was superior in dealing with structured noise such as aliased cardiac cycles and in reproducibility (due to variations in seed positioning across experiments and different laboratories). Structured noise such as aliased cardiac cycles may be relevant in human studies, but because the heart rate of rats (4.5-7.5 Hz) is so much higher than the sample frequency (0.5 Hz) this was not an issue in our experiment. We enhanced the reproducibility of each seed region analysis by automatically positioning the seed in the center of mass of the matched atlas brain regions, which adds the constraint that it is easy to reproduce. Other positions may lead to different results.

### 4.2 The influence of pseudo directionality

Comparing the three graph-theoretical methods, we found the most similarity between the undirected methods (SRCC and RCCA). This suggests that the pseudo direction of the MSRA approach was an important factor. The effect of directionality was highlighted by the comparison of MSRA and SRCC. Both methods relied on the very same seed regions and consequently on the same time-courses. The main difference was the correlation procedure: cross-correlation of two predefined regional time-courses for SRCC (leading to undirected networks) and cross-correlating multiple predefined regional time-courses with each voxel time-course for MSRA (leading to pseudo directed networks). Consequently, the MSRA results were more similar to SRCC than to the RCCA method. The higher resemblance of ICA components to MSRA than to SRCC networks was predominantly induced by the pseudo directionality. Additionally, pseudo directionality enhanced reproducibility, which is evident in the lower variability of the MSRA approach compared to both directed graph-theoretical methods.

The pseudo directionality of the MSRA approach is characterized by fixed seed regions but variable target regions. The location of the fixed seed regions are chosen (hypothesis driven) based on anatomical properties provided by the digital atlas of brain structures; whereas locations of the voxels in the target region are determined by the strength of the connection (data driven) reflecting functional relationships. Thus, the MSRA approach combines hypothesis and data driven as well as anatomical and functional features. This is an important factor that distinguishes our method from all other techniques used to analyze RS connectivity.

### 4.3 Global signal regression and anti-correlation

Although distinguished anti-correlated (i.e. negative correlations) RS networks are reported for humans (Fox et al., 2005), for rats (Liang et al., 2012a; Schwarz et al., 2013) and mice (Sforazzini et al., 2014), we focused on positive correlations only. Negative correlations are difficult to interpret, especially if global regression is used as a preprocessing step. It is known that global regression introduces artificial negative correlations (Murphy et al., 2009), but on the other hand global signal regression leads to more robust RS networks in rodents with higher spatial specificity (Liang et al., 2012a; Liska et al., 2015).

As seen in the histograms of correlation values for the three graph-theoretical methods, both undirected methods are far more influenced by negative correlations. Omitting negative correlations does not shift the histogram, it produces only a cutoff. The histogram of our MSRA approach instead is shifted completely in the range of positive correlation values. Therefore, we conclude that our method is less dependent on false negative correlations introduced by the global regression.

Nevertheless, the MSRA approach is capable of investigating these anti-correlated networks by simply inverting the sign of the initial correlation matrices. The evaluation of these anti-correlated networks is beyond the scope of this study.

### 4.4 Anesthesia

In any animal fMRI studies, anesthesia is an important issue because of its potential side effects on the cardiovascular system and the characteristics of spontaneous neural activity (Grandjean et al., 2014; Nallasamy and Tsao, 2011). The effect of anesthesia on functional connectivity has been investigated by several studies which provide evidence that the connectional architecture of brain networks is preserved at low anesthetic doses (e.g. 1% isoflurane) (Gozzi and Schwarz, 2016; Greicius et al., 2008; Liang et al., 2012b; Vincent et al., 2007) and anesthesia depth should be as low as possible to obtain the major topological features of networks mapped in conscious states. In this study, we controlled anesthesia depth by adjusting to the lowest isoflurane dose while maintaining a constant breathing rate during the fMRI session (see methods). As a result, we detected stable ICA components with remarkable similarity to those described by Becerra et al. (2011) in awake rats.

Even low anesthesia leads to reduced reproducibility and impaired thalamo-cortical connectivity (Liang et al., 2013). While the MSRA approach is characterized by an improved reproducibility across animals (cf. Fig. 9), impaired thalamo-cortical connectivity (represented by enhanced variability of connectivity strength) is evident in all three graph-theoretical methods. Interestingly, the pseudo directionality of the MSRA method revealed increased variability of thalamo-cortical connections for seed regions in the thalamic regions, but, not in the cortical regions.

The thalamus is a highly heterogeneous brain region consisting of small nuclei with functional and anatomical diverse connectivity. It is likely that, placement of seed regions within the thalamus reduces the sensitivity of functional connectivity mapping due to this heterogeneity. The data driven correlations of thalamic voxels to seed regions within the much bigger cortical areas were highly reliable. Consequently, the MSRA approach might be a valid approach for the investigation of thalamo-cortical connectivity e.g. at different states of consciousness.

### 4.5 Paired network based statistics (pNBS)

Comparing graphs of functional brain connectivity at different experimental states usually involves mass-univariate test statistics after which the family-wise error rate (FWE) has to be controlled. Choosing a threshold for the p-values of the test statistics is a balance between sensitivity (i.e. true positive rate) and specificity (true negative rate). Since the generic procedures, such as the false discovery rate (FDR) (Genovese et al., 2002), are highly conservative, they result in low sensitivities (i.e. high false negative rate) and thus may not offer sufficient power.

(Zalesky et al., 2010) presented a powerful method to control the FWE called network based statistics (NBS). An essential part of this method is permutation testing: the group affiliation of each subject is randomly permutated and the statistic of interest is recalculated. This procedure presupposes that the subjects of both groups are independent and exchangeable. This assumption cannot be made in our experimental paradigm, so we adapted the NBS to match the conditions of paired mass-univariate statistics. For this purpose we introduced a control session without stimulation between the two RS measurements.

Repeated measurements of RS connectivity reveal fair to excellent reliability (Du et al., 2015; Zuo and Xing, 2014). Though we cannot postulate the null hypothesis of no difference between the two RS measurements is true for all connections, intra-individual variations should be low. Thus, we use the control session to define a first level threshold p-value that rejects the null hypothesis only for few connections – in this study 1% of all connections of both RS networks. Though usually higher than the conservative FDR (q=0.05) this first level p-value can be used as a measure of variation that should be exceeded by our experimental intervention (i.e. whisker stimulation). From this point of view, the reliability of the repeated intra subject RS measurements is crucial. Lower variability in the control study leads to higher first level thresholds with more potential to detect significant differences due to experimental intervention.

Comparing the three graph-theoretical methods, the MSRA approach gave the highest first level threshold and the most significantly modulated connections. Thus, we conclude that the observed low inter-individual variability also reflects a high intra-individual reliability over repeated measurements.

### 4.6 Resting state connectivity modulation due to whisker stimulation

Using the new MSRA approach, we were able to detect distinct short term RS connectivity modulations due to unilateral whisker stimulation. The modulated network consisted of two subnetworks. One network was comprised of the somatosensory and motor cortex, and the other larger one, was comprised of the thalamus, hypothalamus and amygdala. Most of these structures were activated during whisker stimulation, predominantly those belonging to the sensory-motor network. Most connections, especially those between the thalamic and hypothalamic nuclei and the cortical areas, were diminished during RS. This effect is also present in resting state connectivity after the stimulation period. Such a deactivation of resting state connectivity during task performance has been observed in the default mode network (DMN) (Greicius et al., 2003; Shulman et al., 1997). Li et al. (2012) investigated this specific property of the DMN and observed an enhanced extrinsic connectivity between constituent regions together with decreased intrinsic self-inhibition within these very regions. Paradoxically, the combination of both phenomena leads to the observed deactivation patterns. The authors suggest that this dynamic results in an increase of the DMN’s sensitivity to sensory inputs and may optimize distributed processing during task performance (Li et al., 2012). This principle might be part of RS modulation in general and is still present in RS connectivity shortly after the stimulation period.

Whisking is an important tool rodents use to seek for food or react to a possible thread. To initiate an appropriate behavior, neural processing of information received from whisking involves not only sensorimotor areas of the cortex but also subcortical structures such as hypothalamus (Mogenson et al., 1980). The strongest modulated structures of the subcortical network were the right ventromedial thalamus, the left dorsomedial and the right medial hypothalamus and the right basolateral amygdala.

The contralateral ventromedial thalamus is part of the lemniscal pathway that mediates afferent excitatory projections from whisker to somatosensory cortex (Diamond et al., 2008; Yu et al., 2006). The hypothalamus is generally involved in the regulation of metabolic processes and the autonomic nervous system, the dorsomedial hypothalamus takes part in the regulation of blood pressure and heart rate (Stotz-Potter et al., 1996). Thus, its decreased connectivity might be a response to stress induced by the stimulation. The medial hypothalamus is part of circuitry involved in motivated, i.e. defensive, behaviors, (Canteras, 2002; Swanson, 2000). Lesions in the lateral hypothalamus profoundly impair the ability to orient to stimuli on the contralateral side (Marshall et al., 1971). This deficit is not from motor impairments, but rather a lack of responsiveness to the stimulus, which indicates a direct connection between sensory input and the lateral hypothalamus (Marshall et al., 1971; Northrop et al., 2010).

In contrast to the decreased connectivity between hypothalamus and cortex, the RS connection between hypothalamus and amygdala (mainly basolateral amygdala) was strongly enhanced by whisker stimulation. The lateral amygdala nucleus plays a dominant role in emotional learning and fear conditioning (Pape and Pare, 2010). It receives sensory inputs from the cortex and the thalamus (LeDoux et al., 1990), controls their strength and interferes with the acquisition of fear memory (Ehrlich et al., 2009).

Our results indicate that resting state modulation due to sensory stimulation reflects the impression of a prior sensation and related motor output, but it also involves neuronal circuits known to serve basic processes like fear conditioning and emotional learning initiated by the stimulus.

The biological function of resting state networks is not completely understood, but one hypothesis interprets resting state connectivity as a functional gate that allows the retention of prior information and may influence prospective task-dependent network recruitment and related behavioral output (Deco and Corbetta, 2011). The role of resting state networks relating to human sensorimotor learning and memory consolidation has been described in this context (Albert et al., 2009; Gregory et al., 2014; Mazoyer et al., 2009; Tambini et al., 2010). However, its relevance for emotional learning and conditioning related to sensory stimuli such as touch or pain should be subject for further investigation.

### 4.7 Conclusion

In this study we introduced a powerful new method to analyze resting state functional connectivity. The MSRA approach integrates classical seed based correlation and modern graph-theory, as well as hypothesis and data driven analysis (anatomically chosen seed and functionally correlating target regions). In comparison to two undirected graph-theoretical approaches, it resembles ICA components best and is characterized by its high specificity and reproducibility and less influence from negative correlations. In combination with an adaptation of the network based statistics to paired samples, it promises to be a powerful tool to investigate short term modulations of sensory stimuli related resting state connectivity and ultimately impact our understanding of basic brain functions like fear to higher functions such as learning and memory and consciousness.

## Acknowledgements

The authors thank Sandra Strobelt and Johannes Kaesser for their excellent technical assistance. This work was supported by the BMBF (grant numbers NeuroImpa 01EC1403C, NeuroRad 02NUK034D).

